# Insulin Signaling Functions as a Topological Switch That Couples Aging and Heat Stress Responsiveness

**DOI:** 10.64898/2026.07.08.737295

**Authors:** Manny J. Widuch, Sandra M. Siepka, Richard W Carthew

**Affiliations:** Department of Molecular Biosciences, Northwestern University, Evanston IL; NSF-Simons National Institute for Theory and Mathematics in Biology, Chicago IL

## Abstract

While aging and thermal stress are phenotypically intertwined, their transcriptomic signatures remain paradoxically distinct in *Drosophila*. Here, we demonstrate that this parallel modularity is not an inherent genomic constraint but is actively maintained by Insulin/IGF-1 signaling (IIS). Using transcriptomic analysis, we show that high IIS activity partitions the genome into orthogonal regulatory layers where aging and heat stress function as independent additive inputs. Systemic attenuation of IIS triggers a transition to an integrated architecture. In this state, the genome’s perception of heat stress becomes age-dependent, enabling a hyper-adaptive induction of heat shock proteins in old adults—effectively reversing the typical age-related decline in proteostasis. We propose that the IIS pathway functions as a topological switch, shifting the genome from a modular growth-mode to an integrated maintenance-mode. Thus, dampening the response to heat stress with age is an adaptable regulatory state rather than an inevitable consequence of cellular senescence.

## INTRODUCTION

Aging and an organism’s response to heat stress are inextricably linked. As an organism ages, its ability to manage heat stress declines [1, 2]. Moreover, when an organism experiences a severe heat stress, it accelerates the aging process [3, 4]. Conversely, exposure to repeated sublethal heat stresses causes lifespan extension of an organism, a phenomenon known as hormesis [4–8].

A primary regulatory molecule that links aging and heat stress is the transcription factor Heat Shock Factor (HSF). When an organism is exposed to heat, HSF induces expression of heat shock proteins (HSPs), which act as molecular chaperones to refold proteins damaged by heat, oxidation, and denaturation [9]. HSPs can also facilitate the degradation of damaged proteins via autophagy and the proteasome. Thus, they counteract the accumulation of misfolded and damaged proteins, which causes the cellular dysfunction known as proteotoxicity. Proteotoxicity is a hallmark of aging and environmental stress [10].

Normal aging is associated with increased HSP gene expression [11, 12]. As organisms age, damaged proteins progressively accumulate, triggering elevated expression of HSPs to manage the proteotoxic stress. It is thought that hormesis is at least partially mediated by the upregulation of HSPs after repeated mild heat stress [13, 14]. Interestingly, the induction of HSP expression by acute heat stress is dependent upon an organism’s age. In mammals, the inducibility of HSP expression by HSF declines with age [15–17]. A similar phenomenon is seen in *C. elegans* [18, 19]. Older adults cannot mount a robust HSP response, leading to the accumulation of misfolded protein aggregates. In *Drosophila*, the expression of HSPs in response to heat stress initially increases in middle age before declining in old age [20, 21].

The connection between heat stress sensitivity and aging is further demonstrated by various genetic experiments. Genetic manipulations that increase or decrease HSF levels result in longer or shorter lifespans, respectively [12]. Another transcription factor, FoxO, regulates expression of a subset of HSP genes [22–24]. Mutations that constitutively inhibit or activate FoxO activity result in shorter or longer lifespans, respectively [25–27]. Finally, artificial overexpression of HSP genes leads to greater longevity [9, 11, 28, 29].

HSP genes are not the only ones that are regulated by the process of aging. Approximately 10% of the *Drosophila* transcriptome (∼1,500 genes) changes during aging, and certain classes of genes show strong sensitivity to age [30]. Genes involved in innate immunity, autophagy, and proteolysis are upregulated, while genes involved in oxidative phosphorylation and ATP synthesis are downregulated [31–34]. Acute heat stress also leads to a transcriptome-wide response, with a change in expression of about 1,500 genes in *Drosophila* [35, 36], and a similar number of genes in *C. elegans* [37]. However, of the 1,500 heat-responsive *Drosophila* genes, only 155 of them also vary with the age of unstressed adults, including some of the HSP genes [36].

This observation raises a paradox. If aging and heat stress are phenotypically linked (hormesis, protein damage), why are they transcriptomically distinct? It may simply reflect the central importance of a small subset of genes for both processes. However, the question of why there are so few genes that respond to both has not been answered.

Here, we explore this issue by studying the transcriptome as it responds to age and recovery from acute heat stress. The genomic response is characterized by parallel modularity, where aging and heat stress act as independent, additive layers of genome regulation. However, this additive regulatory network is not a fixed structure but exhibits adaptation in response to the Insulin and IGF-1 signaling (IIS) pathway. The IIS pathway has been long known to affect both aging and responsiveness to stress. Attenuation of the IIS pathway leads to enhanced longevity and increased stress resistance in a broad variety of organisms [4, 38–40].

Remarkably, we find the parallel modularity of the gene regulatory network is not found when the IIS pathway is systemically attenuated. In this state, the regulatory layers collapse into an integrated architecture. Transcriptional responsiveness to heat stress becomes age-specific, with some genes only responding to heat in young adults, and a different set of genes responding to heat in old adults. Overall, it suggests that when IIS activity is reduced, aging and heat stress become interconnected in regulating the organism’s genome at multiple levels. Our findings suggest that the IIS pathway is not simply a volume knob for stress resistance, but is a topological switch for how the genome perceives different stresses.

## RESULTS

It is possible with *Drosophila* to precisely tune the level of IIS activity using genetics. *Drosophila* adults contain 14 insulin producing cells (IPCs) in the brain that secrete the insulin-like peptides dILP-2, dILP-3, and dILP-5 in response to various nutritional cues (Fig. 1A). These dILPs play a major role in promoting growth and reproduction of the animal [41]. One can genetically ablate the IPCs by specifically expressing a pro-apoptosis protein called Reaper (Rpr) in IPCs (Fig 1A). This causes animals to grow and develop to adulthood 70% more slowly even when provided with nutrient-rich food [42]. These IPC-negative adults are smaller, less fertile, but have a longer lifespan and are more stress resistant [42]. We generated such IPC-negative adults for our experiments. We also generated IPC-positive wildtype adults with normal IIS activity (Fig. 1A). Their metabolism is oriented towards growth and reproduction.

**Figure 1.**
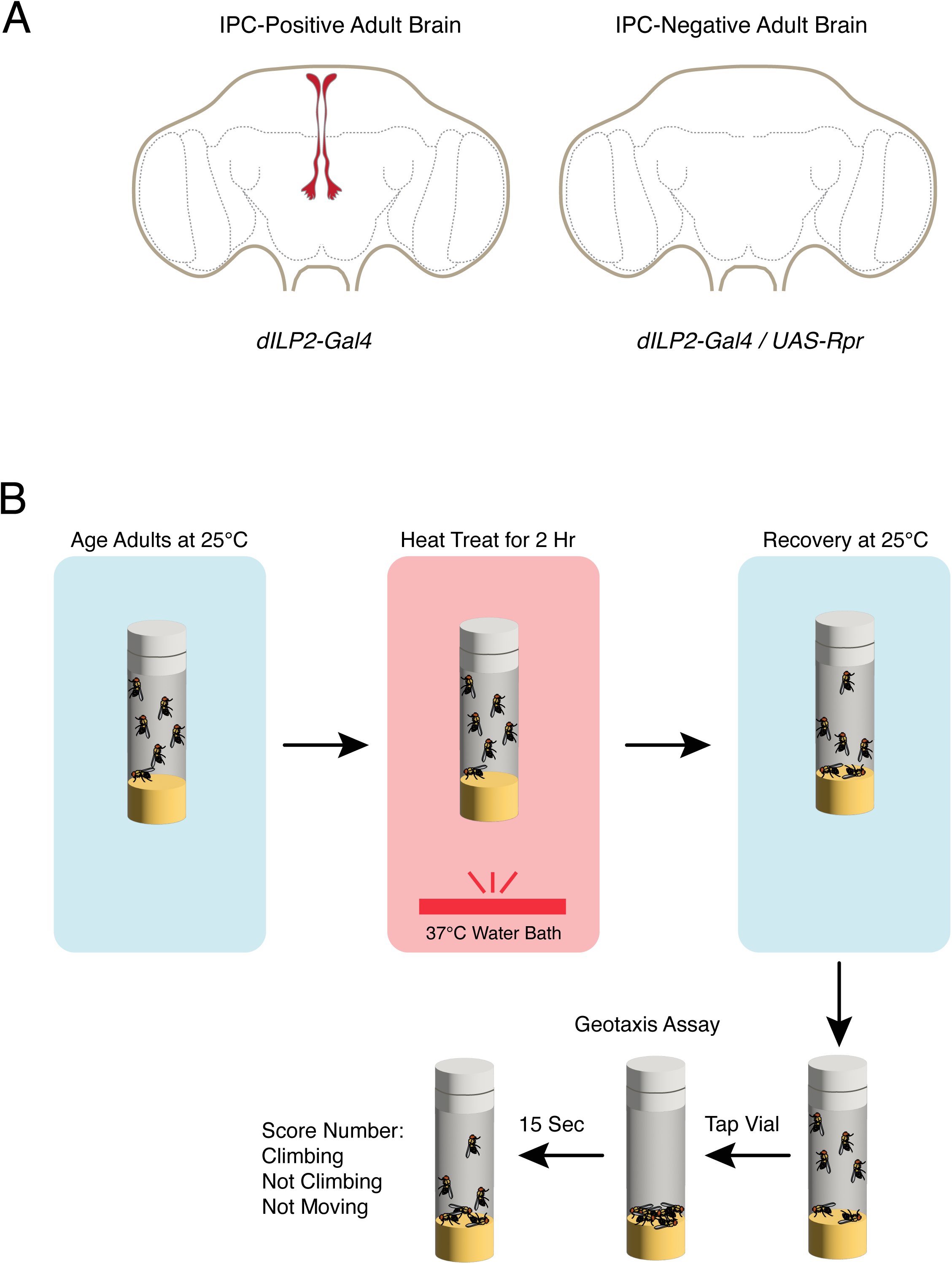
Design of the Experimental Treatments. **(A)** Schematic depiction of the neurosecretory insulin producing cells or IPCs (red) in the adult *Drosophila* brain. They extend processes medially to the tritocerebrum region of the brain as shown, and to the corpora cardiaca and aorta, which are not shown. When an IPC-specific Gal4 driver, dILP2-Gal4, induces expression of the apoptotic protein Rpr, all IPCs are killed (IPC negative). This occurs when dILP2 expression begins in the first-instar larva. **(B)** Workflow of the heat treatment of adults of different ages. Negative geotaxis assays are performed over the timecourse of recovery from heat treatment.

### Locomotor Recovery From Heat Stress Varies With Age and IIS Activity

Aging in *Drosophila* leads to a decline in heat stress resistance [1]. This can be measured in a number of ways including lethality [28]. In young adults (one to five days old), the mortality rate after an acute heat shock (37°C for 60 min) is close to zero. By 40 to 50 days of age, an acute heat shock causes significant mortality.

We wished to know whether any decline in stress resistance with aging could also be detected in the survivors of the heat treatment. Locomotor activity can be measured using the negative geotaxis assay, which exploits an innate behavior [43]. When adults are startled, their instinct is to climb upward away from gravity. This locomotor function naturally declines with age [44, 45]. However, it is unclear how acute heat stress affects climbing and whether this effect is modulated by age.

Therefore, we optimized a heat-stress treatment that was severe enough to cause significant lethality in old adults (30 days old) without killing the entire group. Incubation of 30 day-old adults at 37°C for 120 min was sufficient to induce 60% mortality within 24 hours post-heat treatment. We used this heat treatment to monitor locomotor function in both 30 day-old adults and young adults (four days old).

The ability of non-stressed adults to climb upward in a negative geotaxis assay progressively diminishes with age [44, 45]. We designed our assay so that even old flies that were non-stressed would still perform successfully. Most geotaxis assays set a specific height threshold on the wall for flies to cross [43]. In our assay, flies in a vial were gently tapped and after 15 seconds, we recorded the number of flies that were climbing the wall at any height. Using this relaxed performance criterion, we found that 100% of old flies, who had not been heat-stressed, performed the test successfully.

We subjected old and young adults to a two-hour 37°C heat treatment (Fig. 1B). Only females were assayed, since sex-specific differences have been observed with aging *Drosophila* [46]. Another confounding factor we anticipated was the circadian cycle, which naturally regulates locomotor activity. Moreover, the circadian system interacts with aging and stress resistance [47]. Therefore, we entrained all adults on a 12:12 hr light:dark cycle, and we initiated all heat treatments at ZT 0.

After heat treatment, adults were then allowed to recover at 25°C for 24 hours. Over the course of the 24 hour recovery period, geotaxis assays were performed (Fig. 1B). In addition to counting the number of adults who were climbing on the walls (Climbing Class), we also counted adults who were not climbing but were moving in some manner on the food, whether walking or simply moving a body part (Not Climbing Class). We also counted adults who displayed no detectable movement (Not Moving Class).

When young (four-day old) IPC-positive adults were subjected to the heat treatment, there was an immediate 45% decline in the number of adults who passed the climbing test, followed by a gradual recovery back to a 75% success rate by 24 hours (Fig. 2A). This recovery was accompanied by a reduction in the number of adults who were not climbing, presumably because adults not climbing early in the recovery period regained their ability to climb. About 18% of adults did not move during the recovery period, presumably because they were dead. Accounting for these losses, even after 24 hours, the surviving adults had not fully recovered to their pre-stress locomotor performance level.

**Figure 2.**
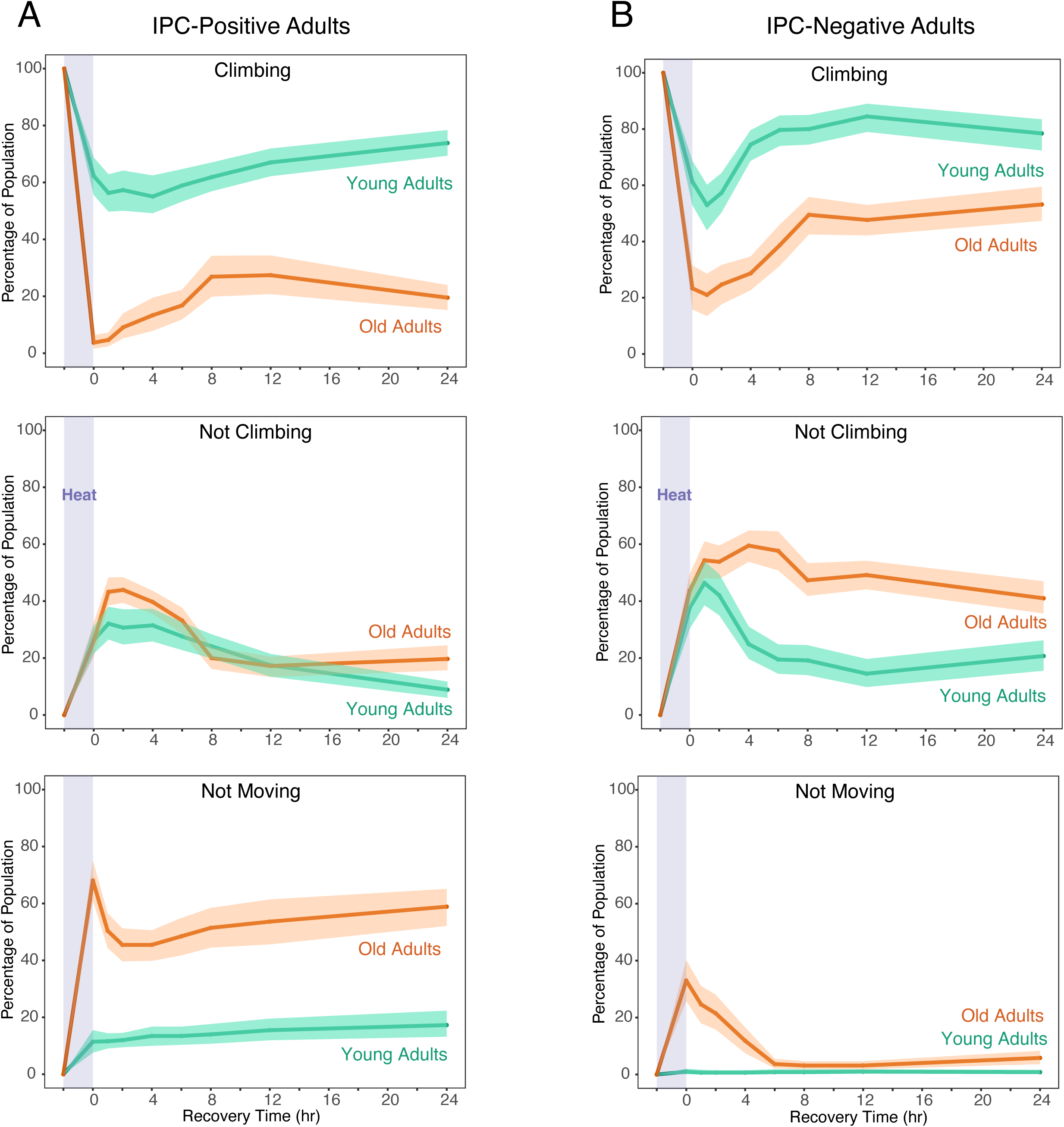
Locomotor Activities in Adults After Heat Treatment. Adults were scored at each timepoint as (1) Climbing, (2) Not Climbing, and (3) Not Moving. The average percentage of the total population who fall into a particular locomotor class is represented by the lines, and the 95% confidence intervals of those estimates of the mean are represented by shading. Young and old adults are distinguished as indicated. **(A)** Locomotor activities in IPC-positive adults. **(B)** Locomotor activities in IPC-negative adults.

Old IPC-positive adults displayed weaker recovery from heat stress (Fig. 2A). Virtually no flies passed the climbing test immediately after heat treatment, and only 20% of adults successfully passed the climbing test after 24 hours of recovery. Immediately after the heat treatment, there was a transient burst in the number of old adults who were not moving, which then decayed to a constant level of 60% of adults who were not moving and presumably dead. Thus, aging of IPC-positive adults leads to impaired recovery in locomotor function and viability after acute heat stress.

We also subjected IPC-negative adults to the heat stress paradigm. Young IPC-negative adults displayed a dip and recovery in their climbing ability that was similar to the young IPC-positive adults we had previously assayed (Fig. 2B). After 24 hours, 80% of young IPC-negative adults were able to successfully climb the walls. Young IPC-negative adults who were not climbing also had similar dynamics with their IPC-positive counterparts. However, the most striking difference between young IPC-negative and IPC-positive adults was in the number of animals who were not moving and presumably dead. There were no dead IPC-negative animals, in contrast to 18% lethality of the young IPC-positive adults.

A different situation was seen with old IPC-negative adults (Fig. 2B). After 24 hours, 50% of old IPC-negative adults were climbing, in contrast to 20% of old IPC-positive adults. A larger number of old IPC-negative adults were moving but not climbing when compared to their IPC-positive counterparts. The reason for this greater recovery of old IPC-negative adults was an almost total lack of lethality after heat stress. Less than 10% of old IPC-negative adults were dead after 24 hours, as compared to 60% of old IPC-positive adults.

Thus, attenuation of IIS activity provided stronger heat stress resistance to old adults. Stress resistance of young adults was less affected by lowered IIS activity.

### RNA-Seq Analysis

To better understand the relationship between aging and the heat stress response, we performed RNA-Seq on the transcriptomes of adults. This was done on young and old adults as they recovered over the first eight hours after heat treatment. We chose this time span because the locomotor activity of adults after heat treatment displayed robust dynamics over the first eight hours of recovery (Fig. 2). Thereafter, a steady-state level in locomotor activity was reached. We sampled RNA from adults 0, 1, 2, 4, 6, and 8 hours after heat treatment. A non-heat treated control group was sampled at the same time as the group at the four-hour time point.

Our behavioral analysis clearly found a significant number of adults dying post-heat treatment. We did not want to include these in our RNA-Seq experiments, and therefore we only collected RNA from adults who passed the geotaxis climbing assay at each time point.

### Heat Stress and Aging Independently Regulate the Genome in IPC-Positive Adults

Together, the transcriptomes of IPC-positive adults of two ages across six time points and one non-heat treated condition yielded many observations in 13,986-dimensional gene expression space. Insights into such high-dimensional data can be obtained by Principal Component Analysis (PCA), which finds genes that contribute most significantly to the variation seen between conditions [48]. Information is measured by the variance found in gene expression, and principal components (PCs) optimally account for the major portion of that variance.

We found the three top PCs (PC1, PC2, PC3) were sufficient to cumulatively explain 54% of the variance in all of the data from IPC-positive adults (Fig. S1A). We therefore used sparse PCA (sPCA) [49] on the RNA-Seq data in order to identify the genes that were most driving the variation explained by PC1 - PC3. A sweep of *L1* penalty values was performed to find the number of genes that accounted for > 90 - 95% of the variance explained by each PC (Fig. S2A). It was found that a sparsity level of 300 genes for each PC was sufficient to capture > 90 - 95% of the variance explained. Critically, the sPCA process does not prevent the same gene from appearing in multiple sparse PCs if it strongly loads onto more than one component.

We plotted the top three PCs against time of recovery from heat treatment for IPC-positive adults. The primary PC (PC1) was found to capture variation in gene expression that correlated with adult age (Fig. 3A). The heat treatment had little to no effect on the genes in this group. In contrast, the secondary PC (PC2) primarily captured variation in gene expression that correlated with heat treatment and not age of the adults (Fig. 3B). The genes in this group responded strongly to heat stress, and the recovery period did not show any relaxation back to the unstressed state. The tertiary PC (PC3) also captured variation in gene expression that correlated with heat treatment rather than age (Fig. 3C). However, this group of genes showed a vigorous relaxation back to an unstressed baseline state over the first four hours of recovery.

**Figure 3.**
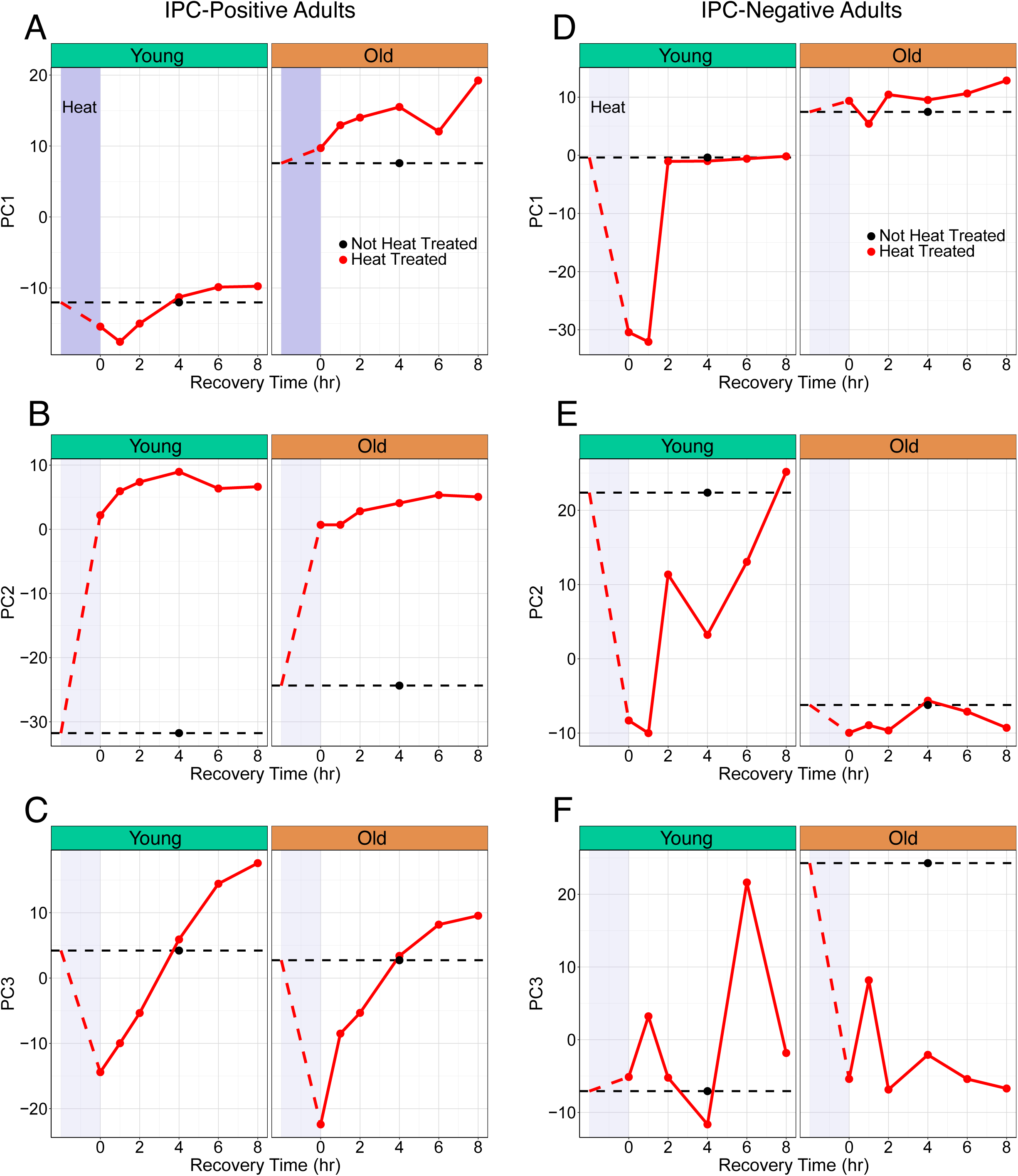
Sparse PCA Analysis Reveals Complex Dynamics. Sparse PCA for each age (young or old) and IIS signaling strength (IPC-positive or IPC-negative), with the coordinates of PC1, PC2, and PC3 for each condition (red circles) plotted against recovery time after heat treatment. The period of heat treatment is indicated by lavender shading. Non-heat treated samples are indicated by black circles. **(A-C)** The time-dependent coordinates of PC1 **(A)**, PC2 **(B)**, and PC3 **(C)** for IPC-positive adults. **(D-F)** The time-dependent coordinates of PC1 **(D)**, PC2 **(E)**, and PC3 **(F)** for IPC-negative adults.

We subjected each group of 300 genes to hierarchical clustering [50]. Each group could be readily classified into two discrete clusters (Fig. S3A-C). We then examined the expression dynamics of all of the genes in each cluster across the time course of stress recovery. The two clusters of PC1 genes exhibited distinct patterns, with one cluster of genes (Cluster A) having greater expression in young adults, and the other cluster (Cluster B) having greater expression in old adults (Fig. 4A,B). Cluster A was significantly enriched for genes involved in oxidative phosphorylation (Fig. S4A). Cluster B was significantly enriched for genes involved in apoptosis, and in Rho1 and p38 stress kinase signaling (Fig S4B). Genes in the unfolded protein response (UPR) and Jun kinase (JNK) pathways were also found in Cluster B (Table S1). These age-related differences had been previously observed [30, 36] and are consistent with the greater metabolic activity found in young adults versus the greater stress activity found in old adults.

**Figure 4.**
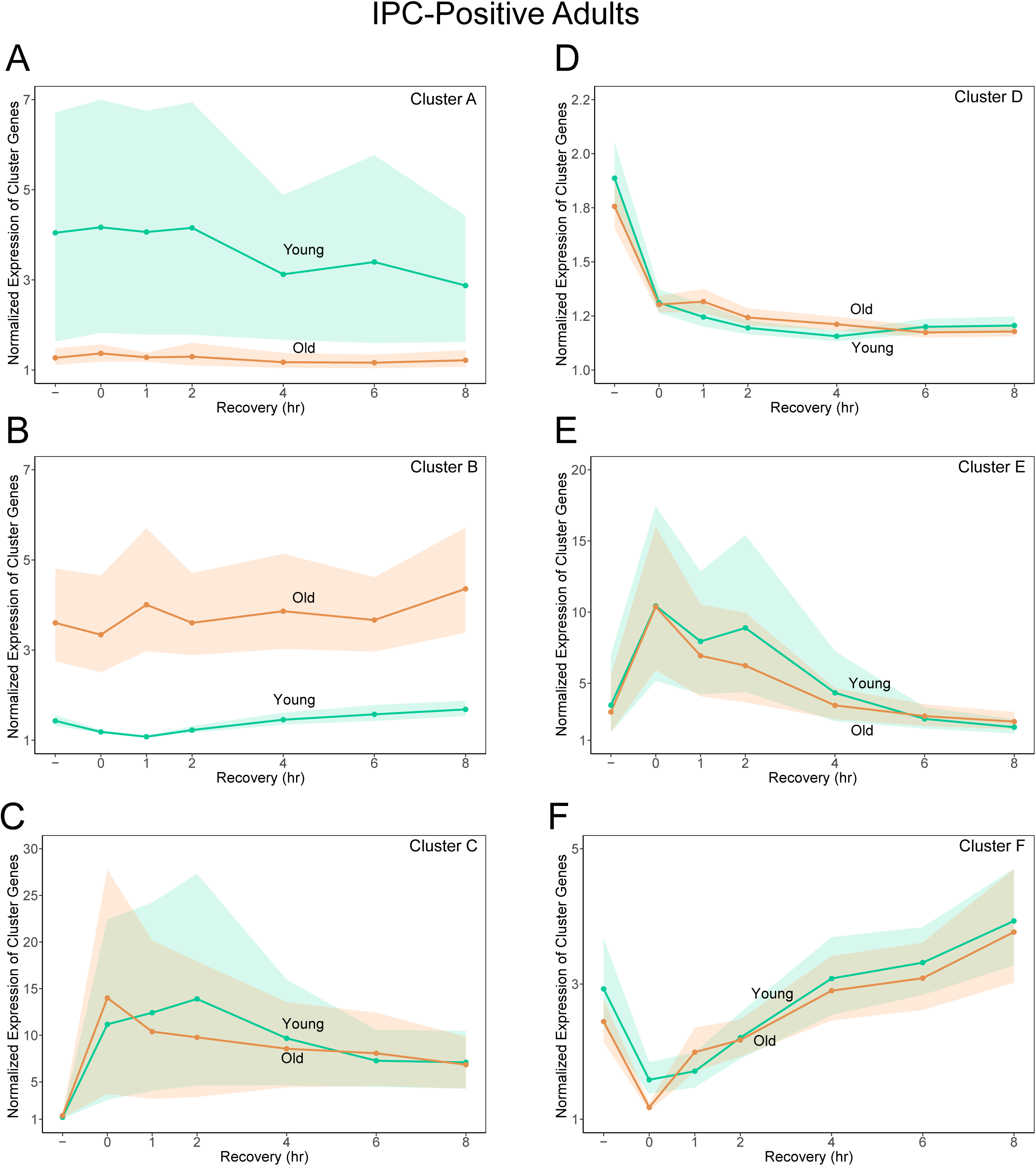
Gene Clusters From IPC-Positive Adults Show Age-or Heat-Dependent Regulation But Not Both. Expression data were normalized on a per-gene basis by dividing each sample’s log-transformed CPM value by the minimum expression value observed for that gene across all samples. This normalization expresses transcript abundance as relative to baseline and allows aggregation of genes within a cluster that have distinct absolute expression levels. Each line represents the bootstrapped estimate of the mean expression value for a given cluster of genes, in young or old IPC-positive adults, plotted against recovery time after heat treatment. Shaded regions represent the 95% confidence levels of those estimates of the mean. **(A)** Cluster A **(B)** Cluster B **(C)** Cluster C **(D)** Cluster D **(E)** Cluster E **(F)** Cluster F.

The two clusters of PC2 genes showed similar expression dynamics in both old and young adults. One cluster (Cluster C) showed strong upregulation of gene expression in response to heat treatment (Fig. 4C), while the other cluster (Cluster D) showed modest downregulation (Fig. 4D). The upregulated Cluster C was significantly enriched for genes encoding HSPs, as well as the translational inhibitor Thor/4EBP (Fig. S4C). Cluster D was enriched for genes involved in rRNA processing (Fig. S4D). The overall response could be interpreted as a heat-induced shift away from protein synthesis and towards refolding of existing proteins.

The expression dynamics of PC3 gene clusters were likewise similar between young and old adults. One cluster (Cluster E) was strongly upregulated by heat treatment but rebounded rapidly (Fig. 4E). Cluster E was enriched for genes encoding HSPs, but a different set of molecular chaperones from those in Cluster C, including the HSPs that are activated by FoxO (Fig. S4E and Table S1). The other cluster (Cluster F) was modestly downregulated by heat treatment and rapidly rebounded as well (Fig. 4F). Cluster F included genes involved in fatty acid metabolism, oxidative phosphorylation, and glutathione-mediated detoxification (Fig. S4F). These are consistent with findings that heat stress induces a decline in fat content [51].

The results of the analyses were striking. The transcriptional response to heat stress was indistinguishable between young and old adults, as evident by the gene expression dynamics of genes associated with PC2 and PC3. Conversely, the primary transcriptional difference between young and old adults was virtually unaffected by heat stress. The results suggested that aging and heat stress have independent and additive effects on the organism’s genome regulation.

One potential limitation of the analysis was that we sequenced only survivors who were climbing. Although this minimized some transcriptomic variation due to physiological variation, sampling was biased for the hardiest survivors, particularly from the old group. The similarity in transcriptomic responses between young and old could have resulted from the fact that only the most resilient old adults were sampled, precisely those capable of mounting a younger-like heat response. Therefore, we also performed RNA-Seq analysis of adults who belonged to the less-resilient “Not Climbing” class.

PCA analysis of this dataset showed similar results to those from the “Climbing” group. PC1 primarily captured genes that were age-responsive and not heat-responsive (Fig. S5A). In contrast, PC2 captured genes with a sustained response to heat treatment in both young and old adults (Fig. S5B). PC3 captured genes with a dynamic response to heat that was similar between young and old adults (Fig. S5C). However in contrast to the “Climbing” group, which showed an immediate and strong response in PC3 to heat treatment, the “Not Climbing” group had a weak initial response followed by a strong overshoot past the baseline unstressed state. Perhaps this difference reflects the different physiological states of the adults. Nevertheless, the RNA-Seq analysis of “Not Climbing” adults validated our finding that the transcriptional response to heat stress is highly similar between young and old IPC-positive adults.

Another potential limitation of the sPCA analysis was the enforced selection of a subset of genes for each principal component by the application of an *L1* penalty. Perhaps there was more overlap between the effects of heat and age if the entire genome was accounted for and not simply 900 genes. We had previously performed standard PCA on all genes in the dataset, not just those selected for sPCA, and we found the top three PCs explained 54% of the variance (Fig. S1A). When the data for all genes was plotted using the top three standard PCs as axes, the 3D plot showed that variation due to age strongly correlated with PC1, while variation due to heat strongly correlated with the other two PCs (Fig. 5A and Supplementary Video 1). The orthogonal distribution of the whole transcriptomic data in the 3D plot supports the conclusion from sPCA analysis that aging and heat stress act as independent regulatory inputs.

**Figure 5.**
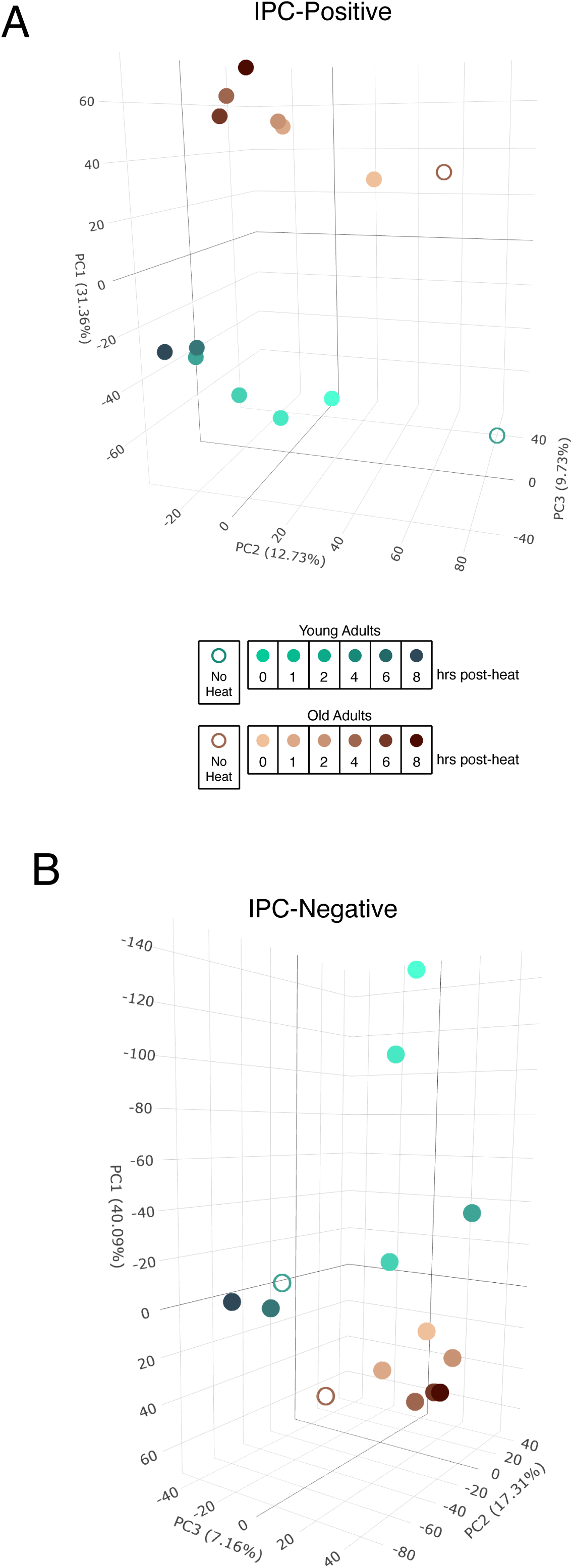
Genome-Wide PCA Analysis of Aging and Heat Stress Recovery. Plots of the complete transcriptomic data for young and old adults without heat treatment or at various times after heat treatment. The first three prinicipal components are used as the coordinate axes. The various conditions of age and heat treatment are color-coded as indicated. **(A)** PCA plot for the IPC-positive adults. **(B)** PCA plot for the IPC-negative adults.

### Heat Stress and Aging Integrate to Co-Regulate the Genome in IPC-Negative Adults

We next performed RNA-Seq analysis on the IPC-negative individuals. Previously, it had been found that attenuating the IIS pathway caused transcriptome-wide changes in expression of genes involved in the FoxO and IIS pathways, as well as autophagy, proteolysis, and apoptosis [52]. Therefore, we first compared the transcriptomes of IPC-positive and IPC-negative adults by PCA (Fig. S6). We also detected genes in the same pathways as Graze et al [52] had found, plus genes involved in RhoA signaling and most of the major paracrine signaling pathways (Fig. S6). Thus, IPC ablation mimics the effect of systemic attenuation of the IIS pathway.

PCA analysis was then performed only on the IPC-negative RNA-Seq samples. We found the three top PCs (PC1, PC2, PC3) were sufficient to cumulatively explain 65% of the variance in the data (Fig. S1B). In contrast, the three top PCs for IPC-positive adults explained 54% of the variance. This increase in captured variance suggests that the attenuation of IIS activity leads to a more coordinated and structured genomic response.

We then performed sPCA on the IPC-negative samples after finding the optimal sparsity level (Fig. S2B), and we plotted the top three PCs against time of recovery from heat treatment. All three PCs captured variation in gene expression that correlated with non-stressed aging (Fig. 3D-F). Remarkably, all three PCs also captured variation in gene expression due to heat treatment but in a manner that was biased by the adults’ age (Fig. 3D-F). PC1 captured a strong heat-induced response in young adults that was transient, rapidly relaxing back to the baseline state after two hours recovery (Fig. 3D). However, there was virtually no heat response in old adults. PC2 also captured a strong response in young adults during the early recovery period, when their expression state shifted very close to the expression state of the old adults (Fig. 3E). In contrast, there was no heat-induced response in old adults (Fig. 3E). PC3 captured a strong response in old adults that was rapidly induced by heat and was sustained over the recovery period (Fig. 3F). Young adults showed a greatly delayed response, with a transient state change six hours after heat treatment before relaxing to baseline by eight hours (Fig. 3F).

We subjected these groups of genes to hierarchical clustering, and again, each PC group could be readily classified into two discrete clusters (Fig. S3D-F). The expression dynamics of each cluster were then analyzed (Fig. 6). One cluster of PC1 genes (Cluster G) showed downregulation of gene expression in young adults, while the other cluster (Cluster H) showed upregulation in young adults after heat treatment (Fig. 6A,B). Old adults showed little heat responsiveness. The downregulated cluster was enriched for genes involved in protein metabolism (Fig. S7A). The upregulated cluster was broadly enriched for genes in the FoxO and IIS pathways, as well as the autophagy pathway (Fig. S7B). The overall response could be interpreted as a heat-induced shift away from protein synthesis and towards turnover of damaged proteins.

**Figure 6.**
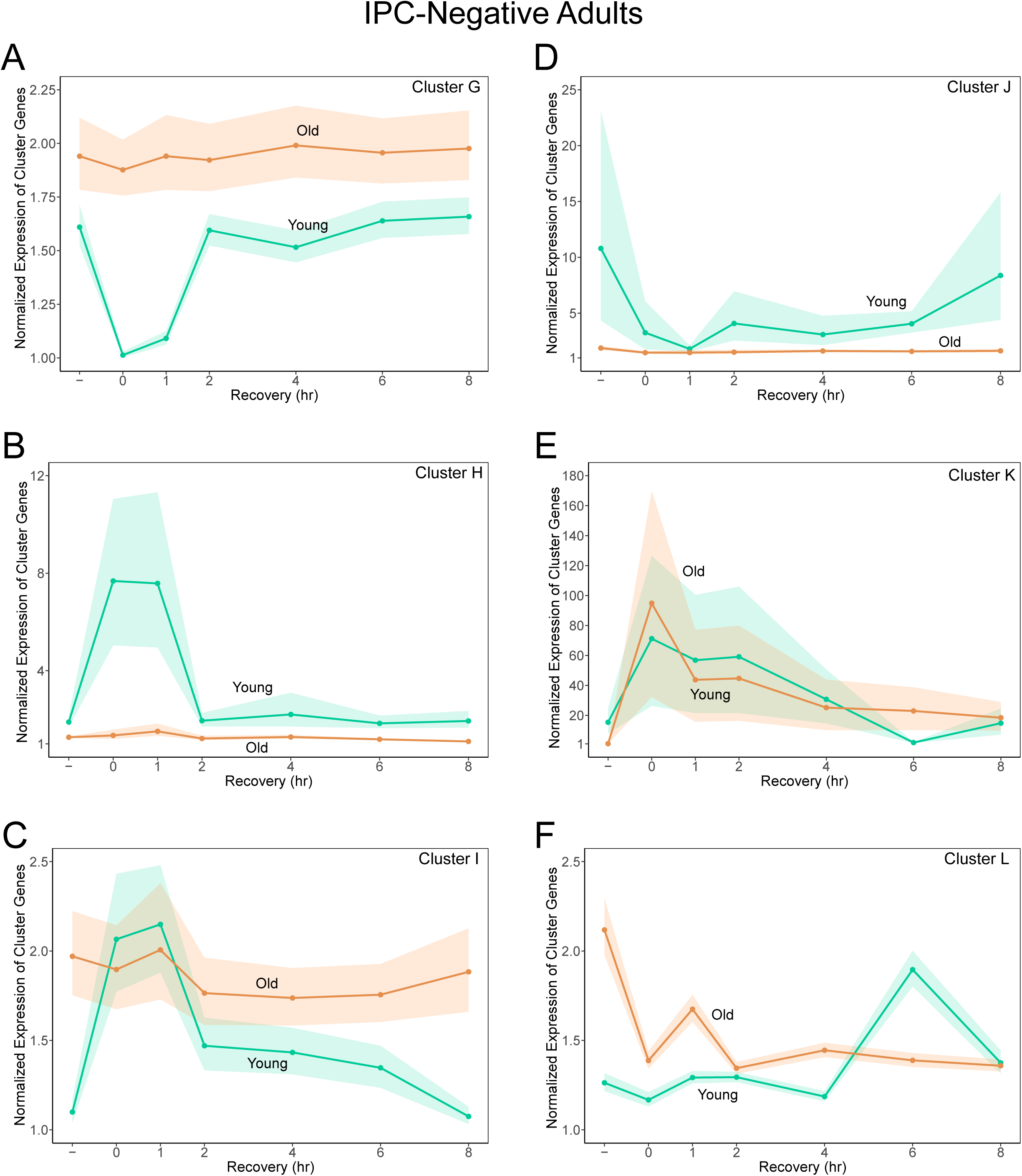
Gene Clusters From IPC-Negative Adults Show Interconnected Age-and Heat-Dependent Regulation. Expression data were normalized on a per-gene basis by dividing each sample’s log-transformed CPM value by the minimum expression value observed for that gene across all samples. This normalization expresses transcript abundance as relative to baseline and allows aggregation of genes within a cluster that have distinct absolute expression levels. Each line represents the bootstrapped estimate of the mean expression value for a given cluster of genes, in young or old IPC-negative adults, plotted against recovery time after heat treatment. Shaded regions represent the 95% confidence levels of those estimates of the mean. **(A)** Cluster G **(B)** Cluster H **(C)** Cluster I **(D)** Cluster J **(E)** Cluster K **(F)** Cluster L.

The expression dynamics of the PC2 clusters were likewise age-biased. One small cluster (Cluster I) was modestly upregulated in young adults by heat treatment (Fig. 6C). This cluster was also expressed at a higher level in old versus young non-stressed adults, and was enriched for genes involved in apoptosis (Fig. S7C). The other cluster (Cluster J) was strongly downregulated in young adults by heat treatment (Fig. 6D). This cluster was expressed at a higher level in young versus old non-stressed adults, and was strongly enriched for genes involved in energy and carbohydrate metabolism (Fig. S7D).

A small cluster of PC3 genes (Cluster K) was expressed at a higher level in unstressed young adults than old adults (Fig. 6E). These genes were transiently upregulated after heat treatment. Remarkably, the peak amplitude of the pulse was 100-fold above baseline in old adults, while it was only five-fold above baseline in young adults. Cluster K was enriched for genes encoding HSPs (Fig. S7E and Table S1). Overall, it suggests that both young and old IPC-negative adults activated the heat shock pathway, but contrary to expectations, there was hyper-responsiveness in old adults. The other PC3 cluster (Cluster L) was expressed at a higher level in unstressed old adults, and was downregulated in old adults after heat treatment for eight hours (Fig. 6F). Young adults showed a delayed response to heat treatment, with a transient upregulation six hours into recovery. Cluster L was enriched for genes involved in cell replication (Fig. S7F).

We had previously performed standard PCA on all genes in the IPC-negative dataset, not just those selected for sPCA, and found the top three PCs explained 65% of the variance. When this multivariate data was plotted with the top three PCs as axes, it showed that age and heat stress correlated with all three PCs in a similar manner as had been elucidated by our sPCA analysis (Fig. 5B and Supplementary Video 2). Thus, the inputs of age and heat are interconnected not just for a subset of genes, but genome-wide.

In summary, the transcriptional response by IPC-negative adults to heat stress was very different from the response by IPC-positive adults. With IPC-negative adults, aging was correlated with heat stress responsiveness. Genes that responded to heat stress were differentially expressed between non-stressed adults of different ages. Moreover, the transcriptional responsiveness to heat stress was age-specific, with some genes only responding to heat in young adults, and a different set of genes strongly responding to heat in old adults. Overall, it suggests that when IIS activity is attenuated, aging and heat stress are interconnected in regulating the organism’s genome at multiple levels.

### HSP Gene Responses Are Modulated by IIS Activity

The heat shock response in *Drosophila* is altered as a function of age. While aging is generally characterized by a decline in proteostasis, the expression of HSPs in response to heat stress often shows a complex, non-linear trend—initially increasing in middle age before declining in old age [20, 21]. To explore this in our RNA-seq data, we compared the raw CPM values for HSP genes between different conditions (Fig. 7). We found that for non-stressed IPC-positive adults, basal HSP gene expression was highly similar between old and young adults (Fig. 7A). Moreover, both old and young adults showed comparable induction of HSP gene expression upon heat treatment (Fig. 7B).

**Figure 7.**
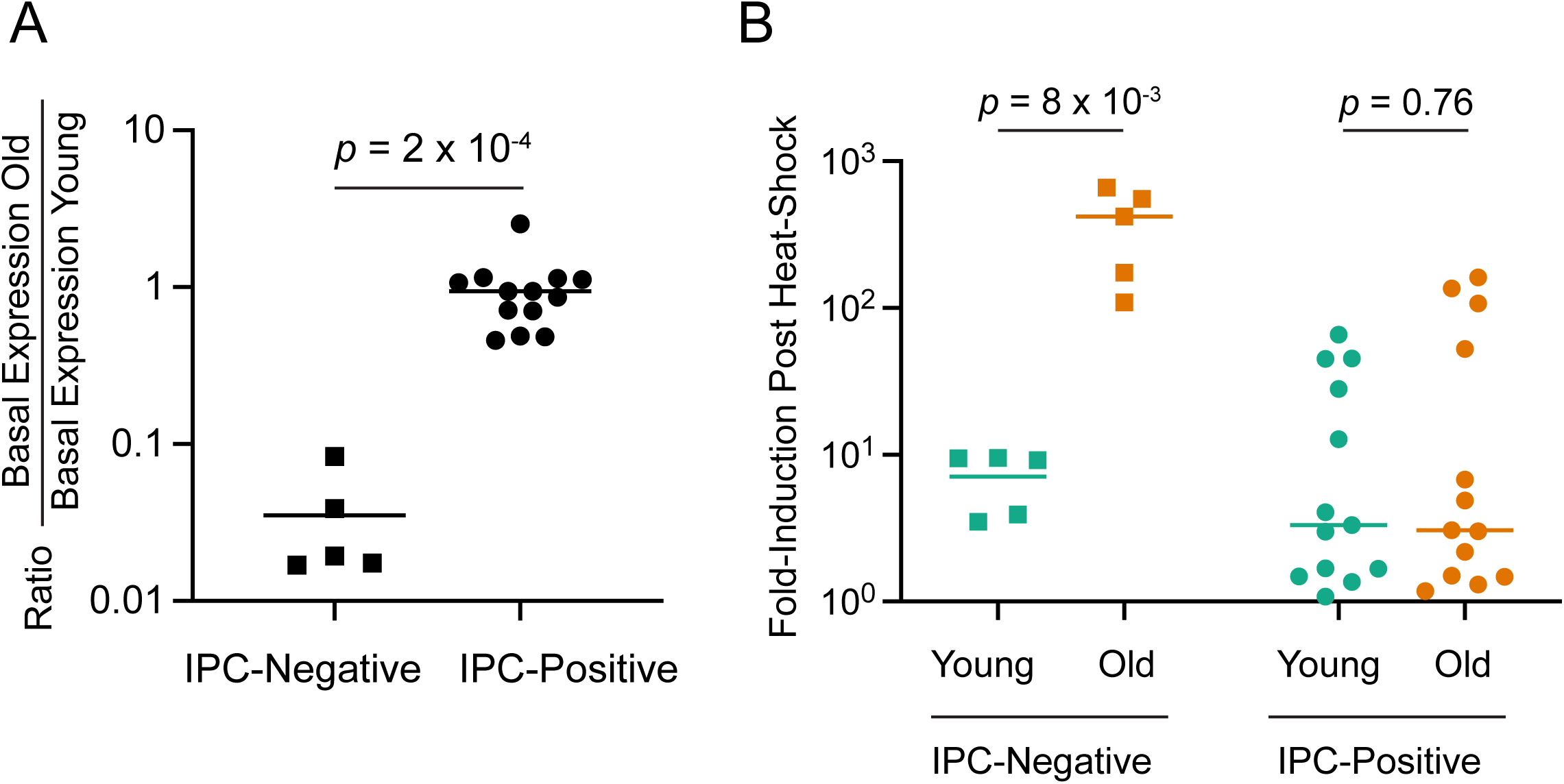
Fold-Change in Expression Levels of HSP Genes. **(A)** The ratio of baseline gene expression of HSP-encoding genes comparing raw CPM levels between old and young adults. HSP genes were identified by sPCA as contributing to variation dependent on either heat stress or aging. Each data point represents a HSP gene expressed in IPC-negative (squares) or IPC-positive (circles) adults. Bars represent the mean. Statistical testing was performed by a two-tailed Mann-Whitney test. **(B)** Fold-increase in gene expression of HSP-encoding genes after heat treatment by comparing raw CPM levels between heat-treated and non-heat-treated adults. These HSP genes were identified by sPCA as contributing to variation dependent on either heat stress or aging. Each data point represents a HSP gene expressed in IPC-negative (squares) or IPC-positive (circles) adults. Bars represent the mean. Statistical testing was performed by a two-tailed Mann-Whitney test.

These trends were greatly altered in IPC-negative adults. Basal HSP expression in the absence of heat-stress was 30-fold lower in old adults relative to young adults (Fig. 7A). However, these old adults had an average 500-fold induction of HSP gene expression upon heat-shock, as compared to an eight-fold induction in young adults (Fig. 7B).

These results focusing on HSP gene expression highlight the hyper-responsiveness of stress genes in old adults when IIS activity is attenuated.

## DISCUSSION

### A Shift in Regulatory Topology: From Parallel to Integrated Architectures

Our findings suggest that the IIS pathway is not simply a volume knob for stress resistance, but is a topological switch for how the genome perceives different stresses. When IIS activity is high, the genomic response to aging and stress is characterized by parallel modularity, where each input generates independent, additive layers of regulation. This is evident by PCA analysis, where age-regulated genes (PC1) and stress-regulated genes (PC2 and PC3) occupy orthogonal axes with minimal interaction. Under this “growth-mode” architecture, the heat stress response remains a fixed-amplitude regulatory program as an organism ages. This fixed program likely declines in its potency as an organism’s proteostatic capacity inevitably diminishes with age, leading to the high lethality observed in old adults after heat treatment.

Conversely, attenuating IIS activity transitions the genome into an integrated maintenance architecture. In this state, the regulatory layers collapse. The primary axes of variance no longer represent isolated inputs but instead capture the interactions between them. This shift is reflected in the significantly higher capture of total biological variance (65%) by the top three PCs, and the coupling of gene expression dynamics to both aging and heat stress. In IPC-negative flies, the heat stress response is no longer an independent module but is instead age-aware. The age of the adult dictates the quality and amplitude of the heat stress response. This integrated topology effectively re-wires the aging fly’s genome, transforming a formerly rigid stress response into a hyper-adaptive mechanism that ensures near-total survival from heat treatment.

The transcriptomic coupling of aging and stress response is also observed when focusing on key genes such as those encoding HSPs. The observed 500-fold induction of HSPs in old IPC-negative adults is a remarkable reversal of the typical age-related decline in proteostatic capacity. Traditionally, the aging *Drosophila* is characterized by a dampening of the HSF response, leaving older individuals vulnerable to protein aggregation and thermal lethality [9]. However, our results demonstrate that this decline is not an irreversible consequence of cellular senescence, but rather a state of active repression maintained by high IIS activity. By attenuating IIS activity, we likely removed a “brake” on the stress response machinery.

How might the IIS pathway re-wire the aging genome? One explanation is that IIS activity simply changes the activity of existing transcription factors. For example, the IIS pathway is known to regulate the transcription factor FoxO [53]. FoxO mediates the effects of the IIS pathway on lifespan in *C. elegans* [25, 26], and it also impacts lifespan and stress resistance in *Drosophila* [27, 54, 55]. When IIS activity is high, FoxO is trapped in the cytoplasm of cells. In this state, the organism focuses on growth and reproduction but has lower stress resistance [53]. When IIS activity is low, FoxO enters the nucleus and activates a wide class of genes involved in stress resistance, including HSP genes and genes required for autophagy, apoptosis, metabolism, and oxidative stress. Perhaps when IIS activity is low, cross-talk between aging and heat stress is enabled by synergistic interactions between the FoxO and HSF proteins at various sites in the genome. However, i-*cis*Target analysis of the genes belonging to Cluster K failed to find binding sites for FoxO in regions with strong HSF binding (data not shown).

Another possible explanation is that IIS activity fundamentally alters chromatin accessibility, allowing transcription factors to bind to different sites in the genome, locking or unlocking genomic regions. Many histone modifications are driven by essential metabolites that are generated by carbohydrate metabolism [56], and epigenetic chromatin modifications occur in aging cells [56]. Metabolomic studies have identified large-scale changes to metabolism in aging *Drosophila* [57]. Perhaps when IIS activity is high, chromatin accessibility to different TFs is partitioned, leading to modular regulation. When IIS activity is low, global epigenetic changes may occur that enable wider accessibility to different TFs mediating stress from aging and heat.

A limitation of our transcriptomic analysis is that the bulk transcriptome measurements are on whole bodies. Gene expression varies with age differently across different tissues and organs [58–60]. Such age-dependent variation can even be seen at single-cell resolution [61]. Therefore, this study is likely focusing on genes whose expression dynamics are consistent across many tissues, making these genes the major drivers of variance captured by our PCA analysis. Future transcriptomic work on isolated tissues and organs will provide a more fine-grained picture of the intersection between aging and stress response as it is programmed by the IIS pathway.

Why is the genome not always regulated by an integrated, hyper-responsive architecture? We propose that parallel modularity represents a growth-mode default, optimized for the rapid ATP synthesis and protein synthesis required for early-life fecundity. In this modular state, the energetic costs of a global, integrated stress response may be prohibitively high. Conversely, the systemic attenuation of IIS triggers a transition to a maintenance-mode integrated architecture. By reallocating the metabolic resources typically reserved for reproduction, the organism enables a global restructuring of the gene regulatory network, granting older adults the hyper-responsiveness necessary to survive proteotoxic stress.

## METHODS

### Drosophila genetics

Two *Drosophila* lines were used: *w; dILP2-GAL4* (on chromosome 3) and *w UAS-Rpr* (BDSC #5823). Individuals ablated of their insulin producing cells (IPCs) were generated by crossing *w; dILP2-GAL4* females to *w UAS-Rpr* males. Only female offspring generated by this cross are IPC-negative owing to their inheritance of the *w UAS-Rpr* X chromosome. *w; dILP2-GAL4/+* females were used as wildtype IPC-positive controls for all experiments. All *Drosophila* were maintained at 25°C on standard cornmeal-molasses food and were entrained to a 12:12 h light:dark cycle.

Virgin females of each genotype (IPC-positive and IPC-negative) were collected. Twenty-five to thirty flies were placed into a vial with standard molasses-cornmeal fly food containing antibiotics (carbenicillin and kanamycin) to limit bacterial growth on the food. Each vial also contained a small dab of yeast paste. Flies that were aged to 30 days were flipped into fresh antibiotic-fortified vials (plus yeast paste) every 4 to 5 days. All vials were incubated at 25°C under a 12:12 h light:dark cycle for the duration of the aging period.

### Heat Treatment

We custom-built a circulating water bath out of a standard non-circulating water bath (30 cm x 32 cm x 13 cm) that we equipped with a culinary Sous Vide machine (Anova Precision Cooker 3.0) placed in one corner of the bath. The water bath thermostat was never turned on, so the Sous Vide apparatus alone was used to heat and circulate the water in the bath for the experiment. The water bath contained 5 plastic 4-way tube racks (DOT Scientific #R1030) such that its capacity was twenty vials. The racks rested atop of six rubber stoppers (size 9) to suspend the racks 2.5 cm from the bottom surface of the water bath. This allowed water circulation around and under the vials, ensuring that the vials were not touching and were evenly distributed in the bath. Water temperature was monitored with a dual probe digital thermometer (FisherScientific #15-078-251). One probe was positioned near the Sous Vide apparatus, the other at the opposite corner of the water bath. Fifteen to thirty minutes before a heat stress was applied, the circulating water bath was preheated to a constant 37.0°C.

The afternoon before heat treatment, aged flies were placed into fresh antibiotic-fortified vials supplemented with a *tiny* dab of yeast paste and were returned to the 25°C incubator. To initiate the heat stress, vials were removed from the 25°C incubator at lights-on time (8 AM) and placed in the 37°C water bath. The vials were weighed down with a 96-position plastic microfuge tube rack to ensure that the vials stayed submerged, and the water surface was level with the midpoint of the vials’ plugs. This ensured that the airspace of every vial was fully submerged, but the water level was not high enough to seep into the vials. After 120 min, the vials were removed from the water bath, gently tapped to ensure that no flies remained stuck to the vial food or yeast paste, and were placed on their sides in the 25°C incubator to monitor recovery.

### Negative Geotaxis Assays

Negative geotaxis is an innate escape response in adult flies. When adults are tapped to the bottom of a vial, they rapidly climb up the vial’s wall. This innate behavior is impaired in locomotor mutants, and it also declines with age. We performed the assay on each replicate vial by tapping it on a table. The flies were individually monitored for their responses, and after 15 seconds, each fly was classified as:

- Climbing: if it was located at any position on the wall of the vial
- Not Climbing: if it was moving in any manner on the food but was not located on the wall
- Not Moving: if it was located on the food, and no limb, head or wing movement was observed

Immediately before each vial was heat treated, the flies were subjected to a geotaxis assay and recorded. Vials of flies were also assayed immediately after heat treatment (removal from the water bath), and at 1, 2, 4, 6, 8, 12 and 24 h after removal from the water bath. Between assay times, vials were kept on their sides in the 25°C incubator on a 12:12 light: dark cycle.

The circadian time at which the heat stress was initiated for all replicates was kept constant at ZT 0. Thus, any effects of the circadian cycle on the geotaxis and transcriptomic responses were the same for all trials. This eliminated any variation in behavioral responses due to variation in timing of the heat stress relative to the circadian cycle.

Each replicate vial of flies subjected to the heat treatment for locomotor assays contained 10 adults. We assayed 58 - 62 replicate vials for each genotype and age for a total of 2,319 flies. Specifically, we assayed 58 replicates of 4d IPC-positive flies (573 flies), 62 replicates of 30d IPC-positive flies (606 flies), 60 replicates of 4d IPC-negative flies (540 flies) and 60 replicates of 30d IPC-negative flies (600 flies). Replicates were not all assayed on the same day but were staggered across multiple days to ensure that results were not due to day-specific batch effects.

For each replicate vial and timepoint, we calculated the percentage of animals that were classified as climbing, not climbing, or not moving. The 95% confidence intervals for point estimates of the mean of the classified data were built by bootstrapping with resampling within each set of replicates grouped by genotype and age. Bootstrapping was repeated 10,000 times. The distributions generated by bootstrapping were checked for normality before obtaining the mean, and the 2.5^th^ and 97.5^th^ percentile values.

### Collection of Samples for RNA-Seq

Flies were aged as described above before undergoing heat treatment. Each replicate vial undergoing heat treatment contained 15 adults. We subjected 20 - 60 replicate vials to heat treatment for each genotype and age, for a total of ∼1,800 flies. Specifically, we heat-treated 20 replicate vials of 4d IPC-positive flies, 60 replicates of 30d IPC-positive flies, 20 replicates of 4d IPC-negative flies and 20 replicates of 30d IPC-negative flies. Replicates of the 30d IPC-positive flies were not all treated on the same day but were staggered across three days.

Immediately after each replicate vial was heat treated, the flies were subjected to a geotaxis assay, and adults in the “Climbing” and “Not Climbing” classes were collected for RNA extraction, as described below. Flies remaining in the replicate vials were then subjected to geotaxis trials at the timepoints: 1, 2, 4, 6, and 8 hours after heat treatment. For each trial, adults in the “Climbing” and “Not Climbing” classes were collected for RNA extraction, as described below. Flies classified as “Not Moving” were not collected since we could not distinguish between dead individuals and living individuals who were paralyzed. Between trials, vials were kept on their sides in the 25°C incubator on a 12:12 light: dark cycle. Flies from replicate vials that were not subjected to heat treatment were collected for RNA extraction from a geotaxis trial at the four-hour time point to serve as a non-heat stress control. This time point was chosen to collect these control samples because it represented the midpoint in the recovery period.

To collect “Climbing” flies from each geotaxis trial, a spatula with the flat end wrapped in double-stick tape was used to capture climbing flies on the wall of a replicate vial. The captured flies were then removed from the spatula with forceps, and placed into a microfuge tube containing 200 µl of TRIzol reagent (Invitrogen 15-596-026). To collect “Not Climbing” flies, those moving on the food of a replicate vial were independently captured with the sticky spatula and placed in a different tube containing 200 µl TRIzol.

As flies were collected at each time point from replicate vials, they were cumulatively added to the TRIzol tube until a total of 10 flies had been added. All 10 flies in a tube shared the same genotype, age, recovery time, and locomotor class. Tubes were stored at-80°C.

### RNA Extraction and Library Preparation

Flies were homogenized using an RNase-DNase free disposable plastic pestle (Kimble Konte K749521-1590). Samples were further homogenized in 800 µL TRIzol reagent by trituration. An aliquot of 500 µL of the homogenate was transferred to a 1.5 mL RNase-free microcentrifuge tube, while the remaining material was stored at −80 °C. Samples were centrifuged at 12,000 × g for 10 min at 4 °C to remove insoluble debris, and the resulting clear supernatant was transferred to a new RNase-free tube and incubated for 5 minutes at room temperature. RNA purification was then performed according to the manufacturer’s protocol for TRIzol reagent, and the RNA was resuspended in 20 µL of RNase free water.

RNA concentration and purity were initially assessed using a NanoDrop One/OneC Microvolume UV–Vis Spectrophotometer (Thermo Fisher Scientific, #ND-ONE-W). For DNase treatment, 10 µg of RNA was diluted in nuclease-free water to a final volume of 80 µL. Samples were treated with 10 µL RQ1 DNase buffer and 10 µL RQ1 DNase (Promega, # M6101), mixed by gentle pipetting, and incubated at 37 °C for 15 min.

RNA was subsequently purified using the Monarch® Spin RNA Cleanup Kit (10 µg; New England Biolabs, Ipswich, MA, USA; Cat. No. T2030) according to the manufacturer’s instructions with minor modifications. These modifications included an additional centrifugation step to ensure complete ethanol removal and a 2-min room-temperature incubation prior to elution spin. RNA was eluted in 30 µL nuclease-free water.

RNA integrity was assessed using Agilent High Sensitivity RNA ScreenTape (Cat. No. 5067-5579), High Sensitivity RNA ScreenTape Sample Buffer (Cat. No. 5067-5580), and High Sensitivity RNA ScreenTape Ladder (Cat. No. 5067-5581) following the manufacturer’s instructions. Samples were analyzed on an Agilent 4150 TapeStation system (G2992AA).

3’ mRNA-Seq was performed on the samples. 3’ mRNA-Seq generates reads biased toward the 3’ end of mRNA transcripts, which has implications for gene-level quantification, particularly for genes with multiple isoforms or complex 3’ UTR structures. However many stress response genes, particularly the HSP genes, encode short transcripts where 3’ bias is less problematic. Libraries were prepared using the QuantSeq 3′ mRNA-Seq Library Prep Kit FWD (Lexogen) with the UDI 12-nt Set A2 (UDI12A_0097-0192; Cat. No. 194.96). During the two bead purification steps, beads were allowed to dry for approximately 8 min prior to elution. Quantitative PCR was performed on a QuantStudio™ 3 Real-Time PCR Instrument (96-Well 0.2ml Block, Thermo Fischer, Cat. No. A28132) to determine the optimal number of cycles required for final PCR amplification. Library quality and fragment size distribution were assessed using Agilent High Sensitivity D1000 ScreenTape (Cat. No. 5067-5584) and High Sensitivity D1000 reagents (Cat. No. 5067-5585) according to the manufacturer’s instructions.

Prepared libraries were sequenced on an Illumina NovaSeq X Plus platform using a 10B flow cell with paired-end 50 bp reads.

### RNA-Seq Read Processing and Alignment

Raw RNA-seq reads were processed for quality control using FastQC (v0.12.0) before and after trimming. Adapter sequences were removed using Trimmomatic (v0.39) in single-end mode using Illumina TruSeq3 adapter clipping (ILLUMINACLIP:2:30:10). Trimmed reads were aligned to the *Drosophila melanogaster* reference genome (dm6; FlyBase release r6.61) using the STAR aligner (v2.7.5a). Alignment was performed using two-pass mapping (--twopassMode Basic) with a maximum mismatch threshold of 2 (--outFilterMismatchNmax 2), and alignments were output as coordinate-sorted BAM files (--outSAMtype BAM SortedByCoordinate). Gene-level counts were generated using the STAR aligner (v2.7.5a) with the --quantMode GeneCounts option in a separate run. During this step, alignments were not written to files (--outSAMtype None). Per-sample count files (ReadsPerGene.out.tab) were aggregated into a gene-by-sample count matrix using a custom AWK script (build_count_table.awk).

Raw gene counts were normalized to counts per million (CPM) to account for differences in sequencing depth across samples. CPM values were calculated by dividing counts for each gene by the total number of reads in the corresponding sample and scaling by 10⁶. Genes with low expression were excluded prior to downstream analysis. Specifically, genes were retained if they exhibited CPM ≥ 1 in at least three samples. The resulting filtered CPM matrix was used for all subsequent analyses.

For variance stabilization, CPM values were log-transformed using log_2_(CPM + 1). Comparison of PCA results using either log-transformed or raw CPM values did not find significant differences in outcomes. For PCA analyses, gene-wise Z-score normalization was applied to the log-transformed CPM matrix, centering each gene to mean zero, and scaling to unit variance. This was done by dividing the center-weighted measures by the standard deviation of the log-transformed CPM data for a given gene and condition.

### Principal Component Analysis

Principal component analysis (PCA) was performed using the prcomp function in R (stats package). The input matrix consisted of Z-score normalized expression values, transposed such that samples were represented as observations and genes as variables.Centering and scaling were disabled (center = FALSE, scale. = FALSE) because the data were already standardized during Z-score normalization.

Variance explained by each principal component was calculated from the eigenvalues, and scree plots were used to assess the contribution of individual components. The first three principal components (PC1–PC3), which accounted for more than 50% of explained variation, were selected for downstream analyses. The scree plots and cumulative variance for each analysis are shown in Fig. S1.

### Sparse Principal Component Analysis (sPCA)

Standard PCA of RNA-seq data often yields dense loading vectors where every gene contributes a non-zero weight, complicating biological interpretation. In contrast, sPCA utilizes an *L1* penalty to enforce sparsity, effectively performing automated feature selection by zeroing out genes that do not significantly contribute to a given component’s variance. sPCA was performed in R using the mixOmics package. Gene-wise **Z**-score normalized expression values were used as input, and the matrix was transposed such that samples were represented as observations and genes as variables. **B**ecause the data had already been standardized, additional centering and scaling were not applied during sPCA (center = FALSE, scale = FALSE).

sPCA was performed across a range of keepX values, to find a threshold value that achieved an optimal balance between sparsity and explained variance. The optimal sparsity level was chosen at a point where additional increases in keepX resulted in minimal gains in variance explained for each component. This corresponded to a sparsity level of 300 genes per component, as shown in Figure S2.

Therefore, the final sPCA model was run with three components and a fixed sparsity parameter of 300 genes per component (keepX = c(300, 300, 300)). Genes with non-zero loadings were extracted for each component, ranked by absolute loading magnitude, and used for downstream clustering and functional enrichment analyses.

### Heatmap Visualization and Gene Clustering

Heatmaps were generated to visualize expression patterns of genes selected by sPCA using the pheatmap package in R. For each principal component, genes with non-zero loadings were extracted and their Z-score normalized expression values were used for clustering and visualization. To determine the number of clusters, the elbow method was applied using the kmeans function from the R stats package. K-means clustering was performed across a range of cluster numbers (k = 1–10), and within-cluster sum of squares was evaluated to identify the point of diminishing returns.

Final gene clustering was performed using hierarchical clustering with Euclidean distance and Ward’s linkage method (ward.D2), implemented using the hclust and cutree functions from the R stats package. The selected number of clusters (k) was used to assign genes to clusters. Prior to heatmap generation, samples were matched to metadata and ordered deterministically based on experimental variables. Samples were first grouped by age, followed by timepoint, and then by sample identifier.

Timepoints were standardized and ordered as: Control, 0, 1, 2, 4, 6, and 8 hours. Control samples corresponded to non-heat-treated samples collected at the 4-hour timepoint and were explicitly labeled as “Control” for visualization.

Heatmaps were generated with gene clustering applied to rows and sample order fixed across columns to preserve temporal structure. For downstream analysis, genes belonging to each cluster were exported for further characterization, including functional enrichment analysis.

### Functional Enrichment Analysis

To determine any functional identity of the gene clusters, genes from each cluster were extracted and subjected to over-representation analysis using the PANGEA (Pathway, Network and Gene-set Enrichment Analysis) tool for *Drosophila* [62]. This tool performs hypergeometric testing using the PypeR function in R. Gene lists from each cluster were analyzed using multiple annotation categories, including: Experimental Data Gene Ontology (Biological Process, Cellular Component, and Molecular Function), pathway databases (DRSC PathON signaling pathway core components and target genes, FlyBase signaling pathways, KEGG, PANTHER, Reactome, and FlyBase metabolic pathways), and protein-level annotations (DRSC COMPLEAT protein complexes and interaction clusters).

Statistical significance was assessed using Benjamini–Hochberg correction, and terms with a false discovery rate (FDR) ≤ 0.1 were considered significant. Filtered terms were used to create gene set node graphs on PANGEA. To validate cluster coherence, we employed a’hit-to-cluster’ ratio as a metric for biological signal strength, comparing the number of genes contributing to a significant GO term against the total cluster membership. Broad, high-level GO categories were used to establish the overall functional density of the clusters; for example, in representative clusters of 277 and 283 genes, the top-ranked categories accounted for 167 and 116 genes, respectively.

To ensure biological precision, these broad signatures were cross-referenced with the most granular GO terms that maintained statistical significance. While granular terms naturally exhibited lower hit-to-cluster ratios, they were used to define the specific physiological identity of the trajectories identified by sPCA. This dual-layered approach confirmed that the clusters were driven by coordinated functional groups rather than stochastic noise.

### Cluster-Based Expression Time Course Analysis

Gene clusters identified from sPCA were used to characterize temporal expression dynamics. A cluster’s gene list was imported and matched to the log-transformed expression matrix (log_2_(CPM + 1), generated using R packages (readr, dplyr, and tidyr), retaining only genes present in the dataset.

To enable comparison across genes with differing expression magnitudes, all expression values were normalized on a per-gene basis by dividing each sample’s expression value by the minimum expression value observed for that gene across all samples. This transformation expresses transcript abundance as relative to baseline and allows aggregation of genes with distinct absolute expression levels.

For each cluster, scaled expression values were used to compute mean expression trajectories across time points. To estimate variability and assess the stability of these trajectories, a bootstrap resampling approach was applied across genes within each cluster. At each iteration, genes were sampled with replacement, and the mean expression across genes in a cluster was calculated for each timepoint. This procedure was repeated 10,000 times, and the resulting distributions were used to compute mean trajectories and 95% confidence intervals using the percentile method. Bootstrapping procedures were implemented using the boot R package.

### Fold-Change Analysis

The raw expression data (CPM) was extracted for genes that had been identified by the sPCA and cluster analyses. The fold-change due to heat treatment was calculated for each gene by dividing the CPM from the 0 hr post-heat sample by the CPM from the non-heat-treated sample. The fold-change due to aging was calculated for each gene by dividing the CPM from the non-heat-treated sample for old adults by the CPM from the non-heat-treated sample for young adults. These fold-changes for specific clusters or groups were either plotted as bar charts with genes as datapoints (Fig. 7). Nonparametric statistical testing of the data employed the Mann-Whitney test.

## Acknowledgements

The Northwestern University NUSeq core facility and Quest high-performance computing facility are acknowledged for providing sequencing and RNA-Seq computational processing services for this work. We acknowledge funding support from the National Institutes of Health (R35GM118144), the National Science Foundation (1764421) and the Simons Foundation (MP-TMPS-00005320).

---

**Figure S1.**
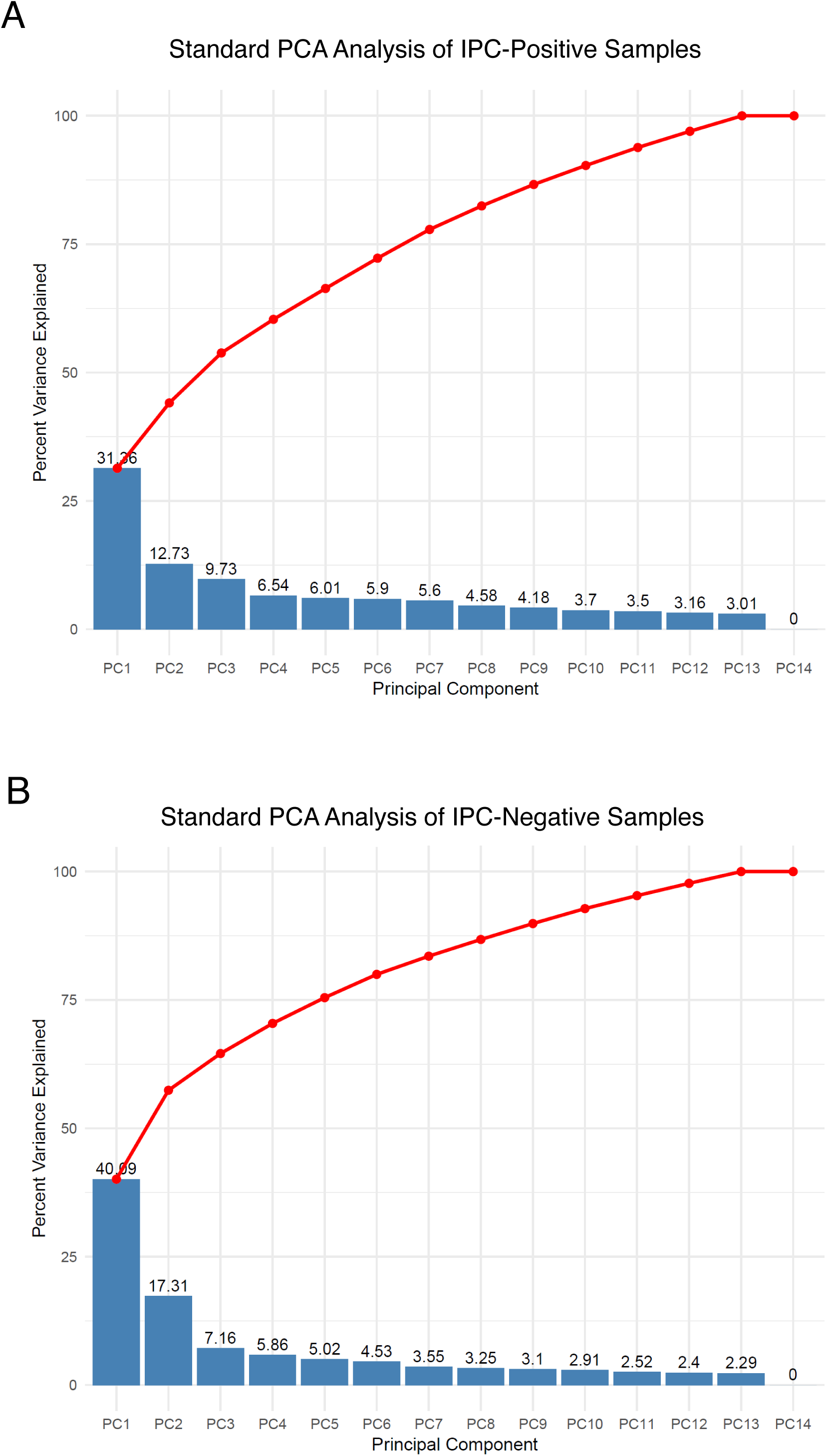
Scree plots of PCA analyses. **(A)** The eigenvalues (variance explained) for each individual component from the IPC-Positive multivariate data. **(B)** The eigenvalues (variance explained) for each individual component from the IPC-Negative multivariate data.

**Figure S2.**
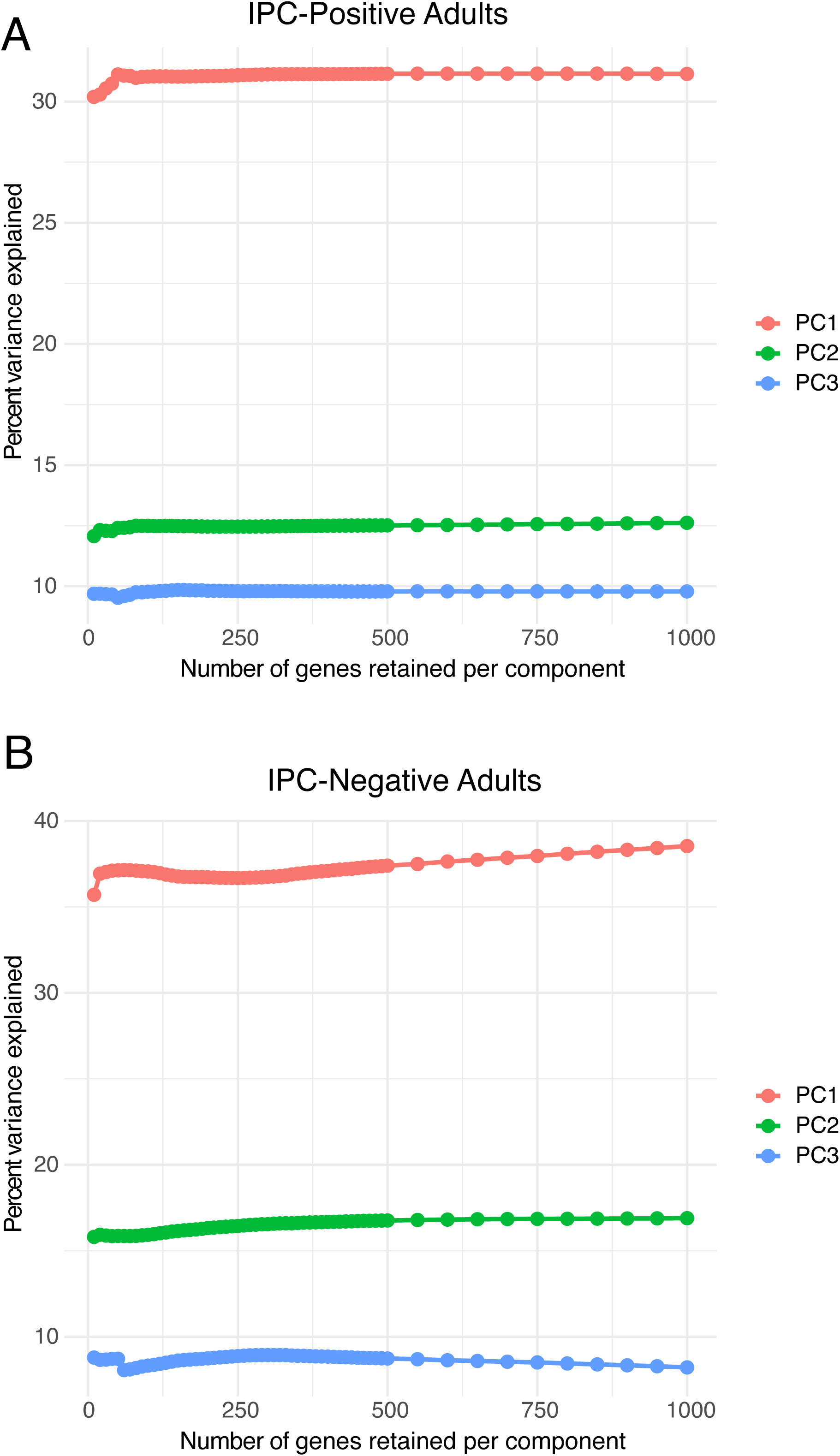
Cumulative Percentage of Explained Variance as a Function of Sparsity. Shown are plots in which sPCA was independently performed using different values of keep(X) for PC1, PC2, and PC3. keep(X) is the parameter defining the number of genes retained. The cumulative percentage of explained variance (CPEV) asymptotes to different levels depending on the PC. The maximal CPEV for each PC is shown in Figure S1. The level of sparsity increases as the number of retained genes decreases. **(A)** The CPEV for the multivariate data derived from IPC-positive adults. **(B)** The CPEV for the multivariate data derived from IPC-negative adults.

**Figure S3.**
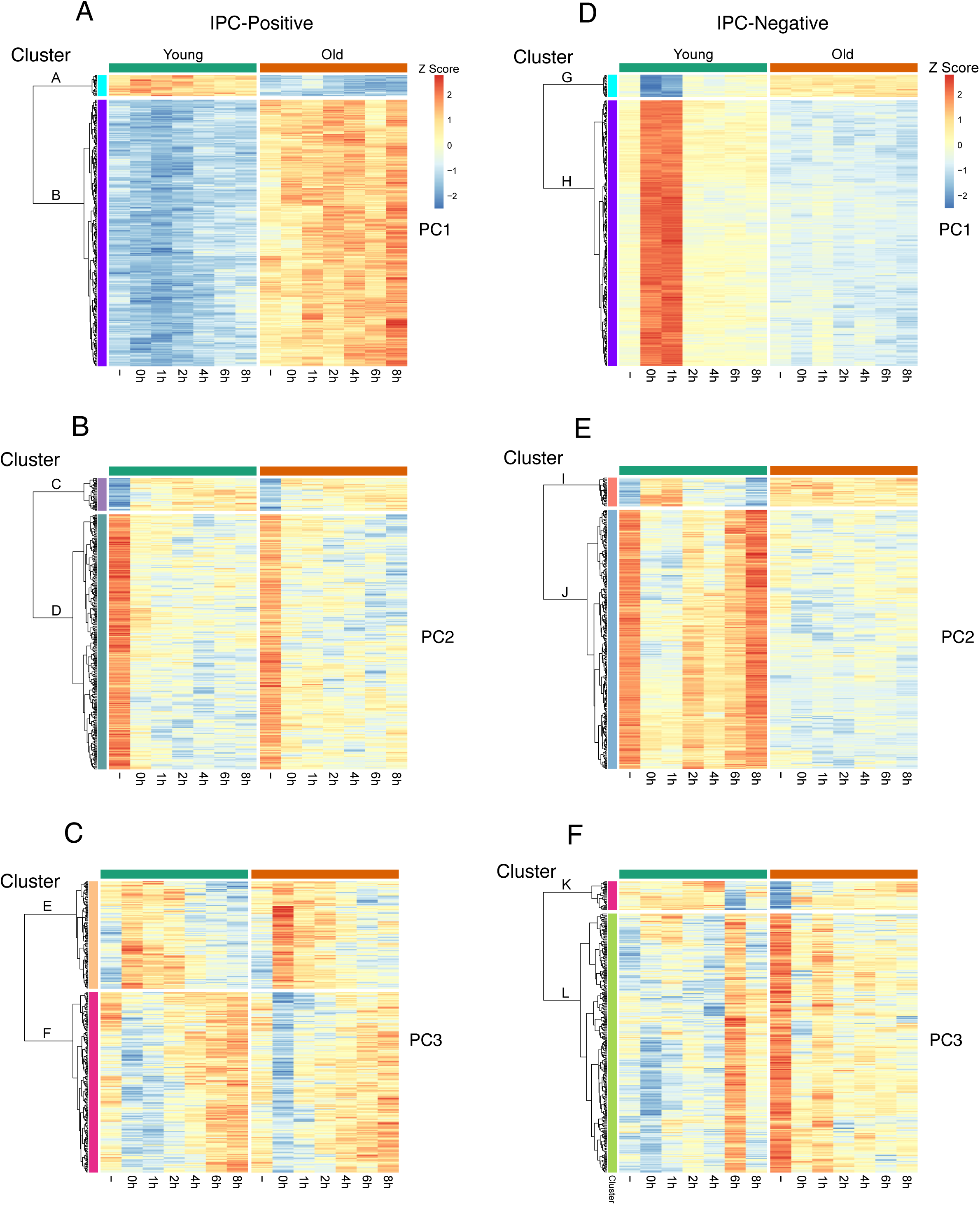
Hierarchical Clustering of Genes. Heatmaps of gene expression, as measured by z-score, for the set of 300 genes (rows) selected for each sparse PC. Columns represent recovery time after heat treatment. The columns marked by a hyphen represent non-heat treated samples. Genes are ordered by their similarity in expression profiles, as determined by hierarchical cluster analysis. Major gene clusters are annotated to the left of each heatmap. **(A-C)** The heatmaps for genes captured by PC1 **(A)**, PC2 **(B)**, and PC3 **(C)** from IPC-positive adults. **(D-F)** The heatmaps for genes captured by PC1 **(D)**, PC2 **(E)**, and PC3 **(F)** from IPC-negative adults.

**Figure S4.**
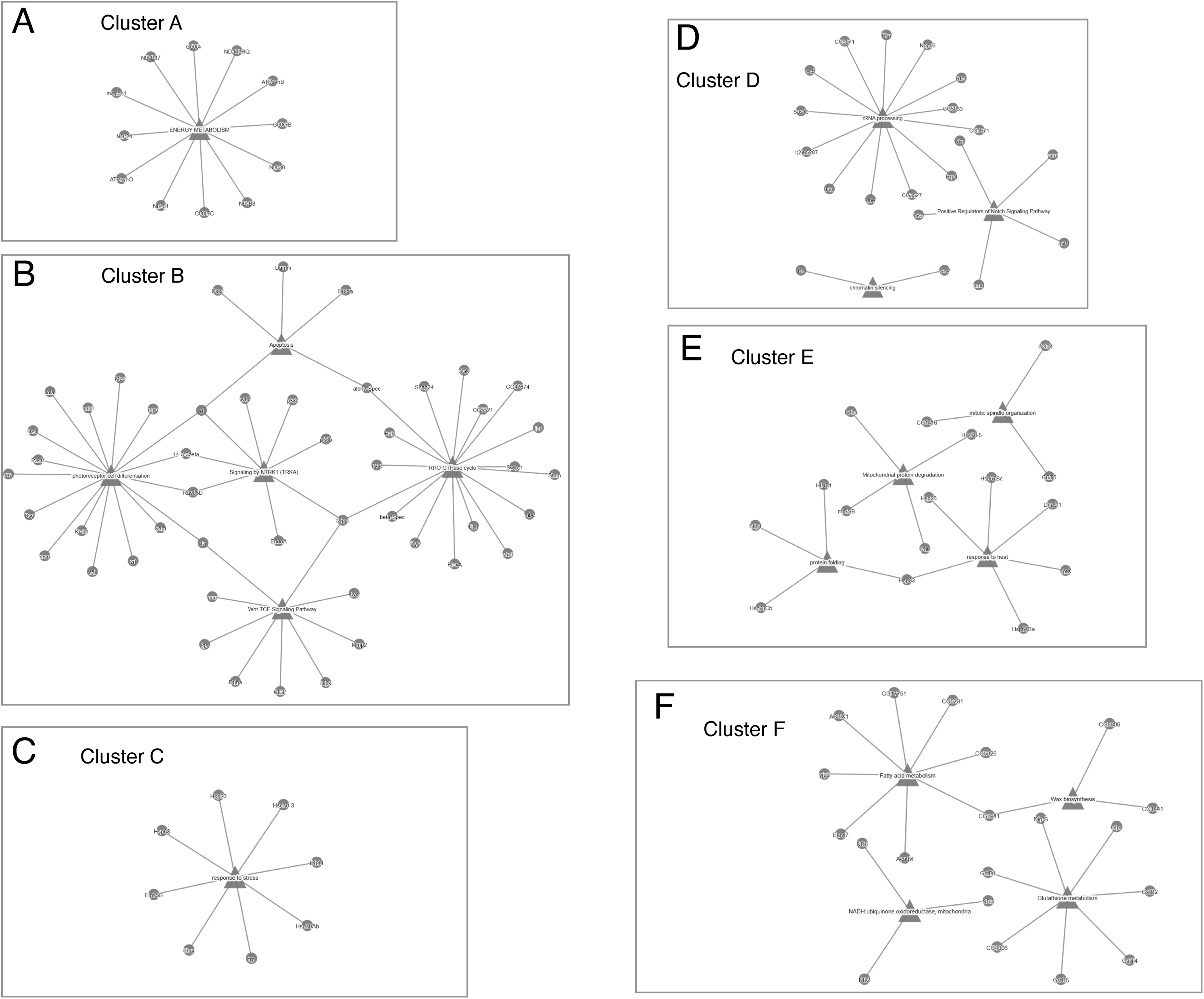
GSEA Analysis of Gene Clusters From IPC-Positive Adults. Genes from each IPC-positive cluster (clusters A to F) were tested for significant enrichment (FDR < 0.1) in a collection of functionally annotated *Drosophila* gene sets. Shown are network graphs of the selected gene sets. Triangle nodes represent the names of the gene sets, and circle nodes represent the genes associated with the gene sets. Edges represent gene to gene-set associations. **(A)** Cluster A **(B)** Cluster B **(C)** Cluster C **(D)** Cluster D **(E)** Cluster E **(F)** Cluster F.

**Figure S5.**
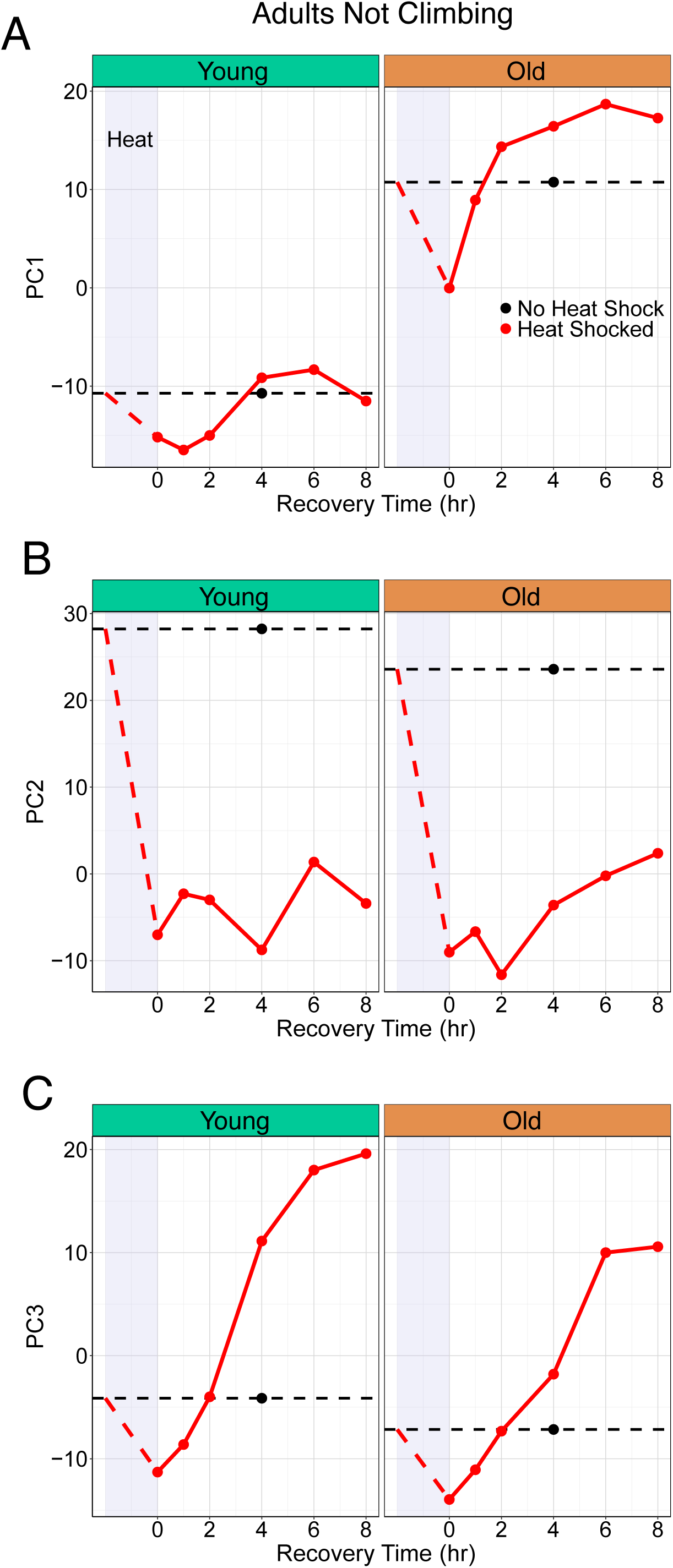
Sparse PCA Analysis of IPC-Positive Adults Classified as Not Climbing in Locomotor Assays. Adults classified as “Not Climbing” were sampled for RNA-Seq analysis after heat treatment. Sparse PCA of this transcriptomic data for each age (young or old) is shown, with the coordinates of PC1, PC2, and PC3 for each condition (red circles) plotted against recovery time after heat treatment. The period of heat treatment is indicated by lavender shading. Non-heat treated samples are indicated by black circles. **(A)** The time-dependent coordinate of PC1. **(B)** The time-dependent coordinate of PC2. **(C)** The time-dependent coordinate of PC3.

**Figure S6.**
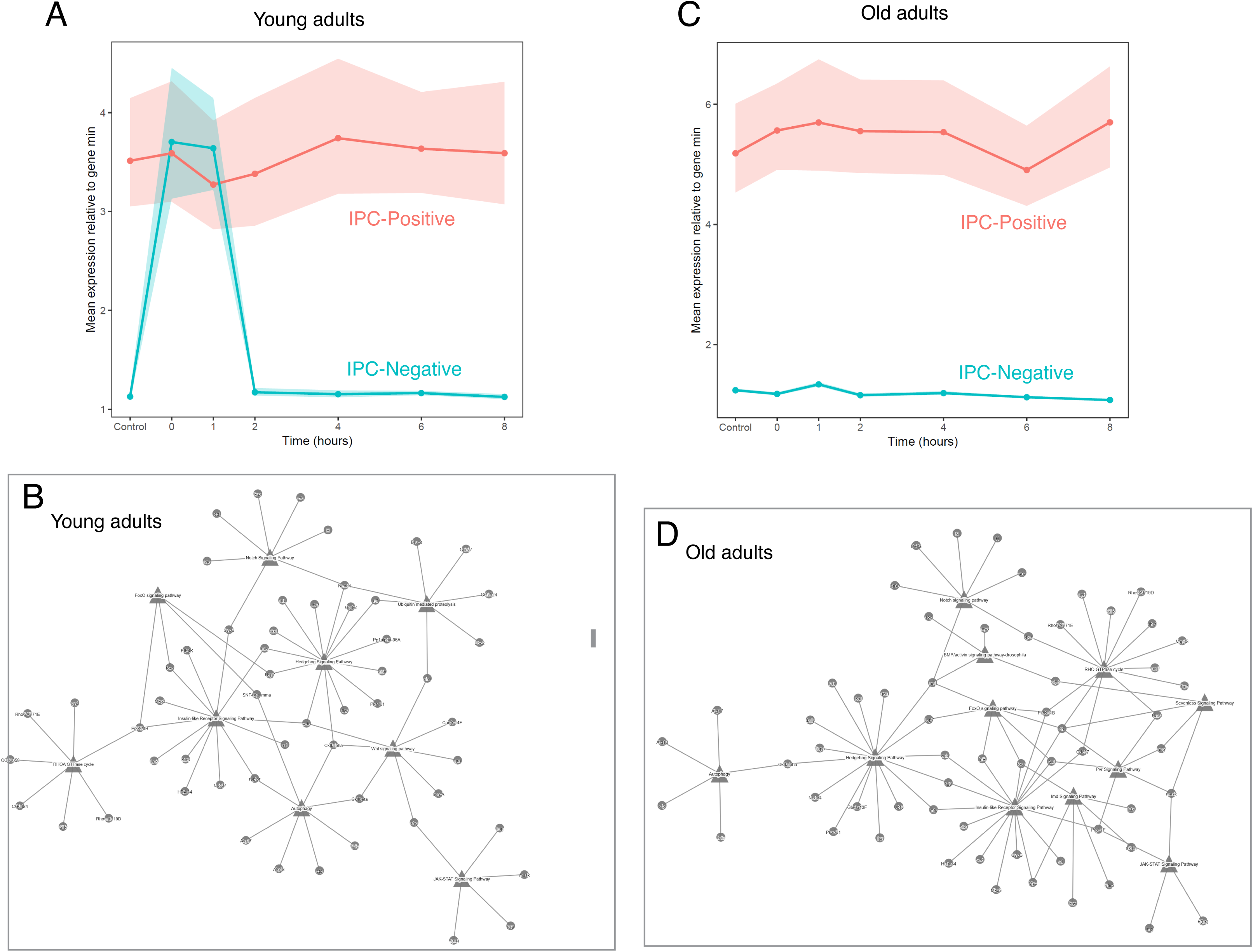
Transcriptomic Comparison Between IPC-Positive and IPC-Negative Adults. **(A)** In young adults, the major cluster of sPC1 genes (298 genes) is differentially expressed between IPC-positive and IPC-negative adults. Expression is higher in IPC-positive adults, except for one hour after heat treatment, when the expression levels are similar between IPC-positive and-negative adults. **(B)** GSEA analysis of the gene cluster shown in (A). The cluster is enriched for genes involved in proteolysis, autophagy, and a variety of signaling pathways. **(C)** In old adults, the major cluster of sPC1 genes (294 genes) is differentially expressed between IPC-positive and IPC-negative adults. Expression is higher in IPC-positive adults. **(D)** GSEA analysis of the gene cluster shown in (C). The cluster is enriched for genes involved in similar functions as found for the young samples — autophagy and many signaling pathways.

**Figure S7.**
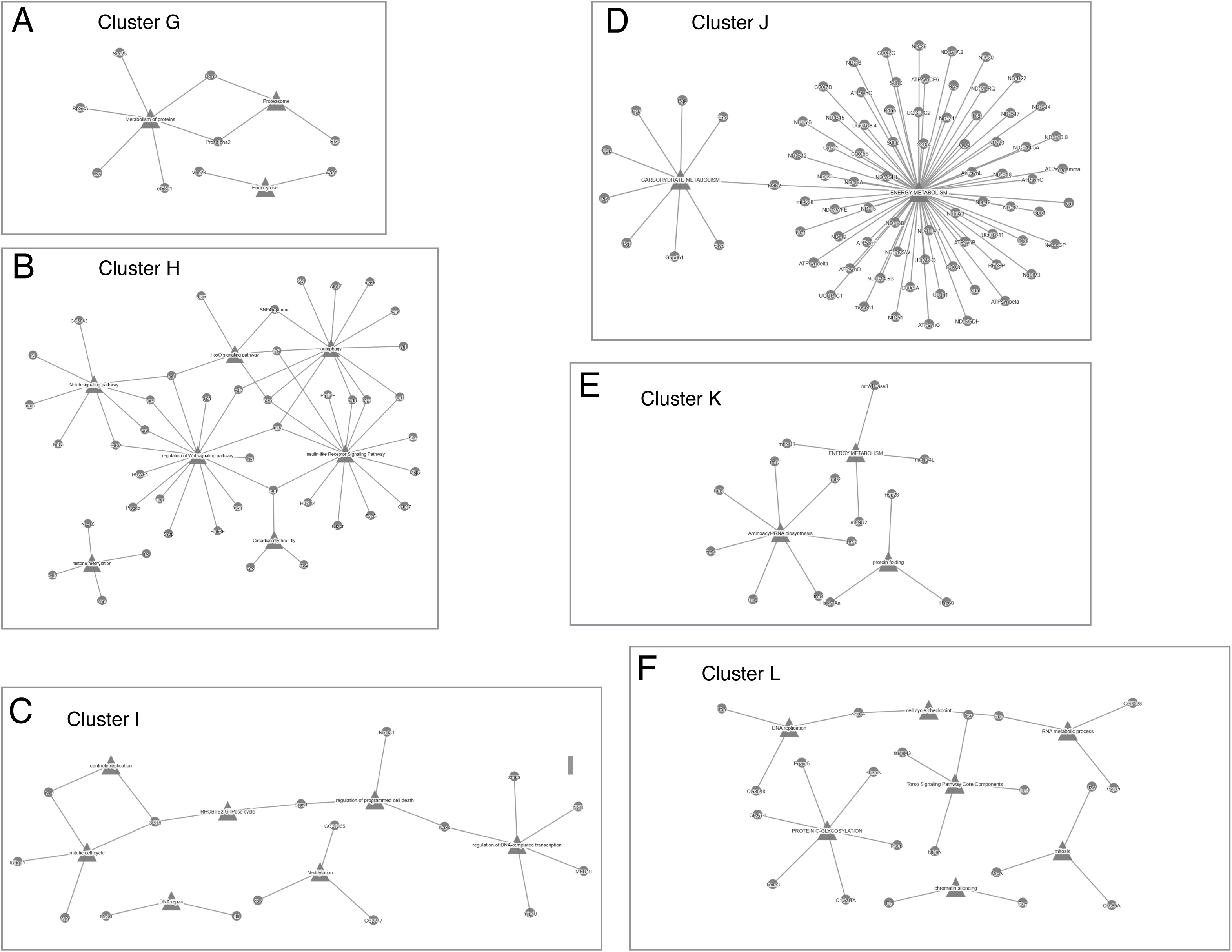
GSEA Analysis of Cluster Genes From IPC-Negative Adults. Genes from each IPC-negative cluster (clusters G to L) were tested for significant enrichment (FDR < 0.1) in a collection of functionally annotated *Drosophila* gene sets. Shown are network graphs of the selected gene sets. Triangle nodes represent the names of the gene sets, and circle nodes represent the genes associated with the gene sets. Edges represent gene to gene-set associations. **(A)** Cluster G **(B)** Cluster H **(C)** Cluster I **(D)** Cluster J **(E)** Cluster K **(F)** Cluster L.

**Table S1.** Lists of all genes that belong in Clusters A to L.

**Supplementary Video 1.** Animation of the 3D PCA plot from Figure 5A.

**Supplementary Video 2.** Animation of the 3D PCA plot from Figure 5B.

## Notes

### Competing Interest Statement

The authors have declared no competing interest.

